# Ca^2+^ increases cardiac muscle viscoelasticity independent of active force development

**DOI:** 10.1101/2025.02.26.640328

**Authors:** Anthony J. Baker, On Yeung Li, Filip Ježek, Paul C. Simpson, Naomi C. Chesler, Daniel A. Beard

## Abstract

In addition to activation of muscle contraction by Ca^2+^, recent studies suggest that Ca^2+^ also affects muscle passive mechanical properties. The goal of this study was to determine if Ca^2+^ regulates the stiffness of cardiac muscle, independent of active contraction. The mechanical response to stretch for mouse demembranated cardiac trabeculae was probed at different Ca^2+^ levels after eliminating active contraction using a combination of two myosin ATPase inhibitors: *para*-nitroblebbistatin (PNB, 50 μM), plus mavacampten (Mava, 50 μM). Myocardial force level was assessed during large stretches (≈ 20% initial muscle length) with a range of stretch velocities. For relaxed muscle, in response to stretch, muscle force rose to a peak and then decayed toward a lower steady-state level, consistent with the viscoelastic nature of cardiac muscle. Peak force was higher with faster stretch velocity, but the steady-state force was independent of stretch velocity, consistent with the presence of both apparent viscous and elastic components of the stretch response. In the presence of the inhibitors PNB plus Mava, when Ca^2+^ level was increased, active contraction was completely prevented. However, the viscoelastic force response to stretch was markedly increased by high Ca^2+^ and was > 6-fold higher than at low Ca^2+^ level. The relationship of viscous force to Ca^2+^ level had a similar form to the relationship of active force to Ca^2+^ (measured in the absence of inhibitors), suggesting a common regulatory mechanism is involved. As expected, Ca^2+^-activated contraction was inhibited by lowering the temperature from 21°C to 10°C. In contrast, the Ca^2+^-activated viscous property was not inhibited at lower temperature, further suggesting that active contraction and the viscous property involve distinct mechanisms. This study demonstrates that in addition to triggering activation of contraction, Ca^2+^ also increases the apparent viscous property of cardiac muscle.

**New and Noteworthy:** Ca^2+^ is well-known to trigger activation of muscle contraction. This study demonstrates a new mechanical role for Ca^2+^ in cardiac muscle involving a >6-fold increase in the apparent muscle viscoelasticity. Activation of a viscous element by Ca^2+^ might influence the mechanical properties of activated cardiac muscle.

## INTRODUCTION

Electrical stimulation of frog skeletal muscle was recently reported to increase the resistance of muscle sarcomeres to stretch, independent of active force development (1). This phenomenon was attributed to the effects of muscle activation, likely due to increased intracellular level Ca^2+^, on the mechanical properties of titin and on titin’s interactions with other sarcomeric proteins. Given the importance of cardiac muscle mechanical properties in health and disease (2, 3), the goal of this study was to determine if and how passive mechanical properties of cardiac muscle are sensitive to levels of Ca^2+^. Using demembranated cardiac trabeculae from male and female mice, we measured muscle force during and following muscle stretches that were imposed with a range of velocities. To measure the effect of Ca^2+^ in the absence of force production, we prevented cross-bridge cycling using a combination of two inhibitors *para*-nitroblebbistatin (PNB) and mavacampten (4, 5). The combination of these inhibitors prevented Ca^2+^-activated force development. As previously reported, we found that the dynamic force response of cardiac muscle to stretch consists of a velocity-sensitive viscous component and a velocity insensitive elastic component (6-9). Importantly, at the fastest stretch velocity, we found that the viscous component of the force response was > 6-fold higher at high Ca^2+^ level compared at low Ca^2+^ level. In summary, for cardiac muscle, activation by Ca^2+^ caused a large increase in a muscle viscous property that was independent of active force development.

## METHODS

The study was approved by the Animal Care and Use Subcommittee of the San Francisco Veterans Affairs Medical Center and conformed to the *Guide for the Care and Use of Laboratory Animals* published by the National Institutes of Health (Revised 2011). This institution is accredited by the American Association for the Accreditation of Laboratory Animal Care (Institutional PHS Assurance Number is A3476-01).

### Demembranated right ventricular (RV) trabeculae

Trabeculae were prepared as we recently described (10) using 12-week old male and female C57BL/6J mice (Jackson Labs). Briefly, hearts were removed from deeply anesthetized mice (3% isoflurane), placed in cold arrest solution and flushed with a modified Krebs solution (10). A piece of the RV near the tricuspid valve that contained a trabecula was dissected and immersed in ice cold relaxing solution (see below) plus 2% Triton X-100 (Sigma-Aldrich) for 1 hour, washed in ice-cold relaxing solution for 1 hour, then stored at -20**°**C for up to 85 days (mean 24 days) in a 1:1 mixture of relaxing solution and glycerol (11, 12).

### Solutions

Ca^2+^-free relaxing solution (pCa 11) contained (in mM): EGTA 20; MgATP 8; creatine phosphate 12; N,N-bis[2-hydroxyethyl]2-aminoethane sulfonic acid (BES) 100; pH adjusted to 7.1 with KOH, ionic strength adjusted to 200 mM with KCl, and temperature 21**°**C (13). Preactivating solution was identical but with calcium-buffering reduced by replacing 19.5 mM of EGTA with HDTA (hexamethylenediamine-N,N,NV,NV-tetraacetate) (Fluka). Activating solution (pCa 4.51) contained 20 mM Ca^2+^EGTA. Relaxing and activating solutions were mixed to obtain solutions with intermediate pCa (14). All solutions contained 1% (v/v) Protease Inhibitor Cocktail P-8340 and 10 IU/mL creatine kinase (Sigma, St. Louis, MO).

To inhibit myosin cross-bridges a combination of 50 μM *para*-nitroblebbistatin (PNB) plus 50 μM Mavacampten (Mava) was added from 10 mM stock solutions dissolved in DMSO (1% DMSO final).

### Myofilament extraction

Treatment of cardiac trabeculae with a high salt extraction protocol removes >95% of myosin and actin which anchor titin in the sarcomere (15). After extraction, non-myofilament structures, especially collagen, the mechanical properties of muscle should be determined primarily by the collagen extracellular matrix. After mechanical studies, trabeculae were incubated with relaxing solution containing 0.6 M KCl for one hour followed by relaxing solution containing 1 M KI for one hour at 21**°**C (15). Muscles were washed in relaxing solution for 60 s and mechanical tests were repeated.

### Mechanical studies

A demembranated trabecula (one per mouse) was attached using aluminum t-clips to a force transducer (Model 400, Aurora Scientific, Inc., Ontario, Canada) and a computer-controlled servo-motor in a small glass-bottomed chamber of a Permeabilized Fiber Test System (Model 1400A, Aurora Scientific, Inc., Ontario, Canada) on an inverted microscope with a video system (Model 900B, Aurora Scientific, Inc., Ontario, Canada) to measure sarcomere length using a 40X objective. Temperature was set to 21**°**C for mechanical studies. The effect of lowered temperature (10**°**C) was also determined.

In relaxing solution, the initial muscle length (Lo) was adjusted to set the sarcomere length to 2.0 µm. Trabecula dimensions were measured and used to normalize muscle force to the muscle cross-sectional area (assuming an elliptical cross-section). Then, maximal Ca^2+^-activated force (Fmax) was measured by moving the trabecula to pre-activating solution for 60s, activating solution for 6s, and then returned to relaxing solution.

In relaxing, activating, and intermediate pCa solutions, and in the presence of 50 μM PNB plus 50 μM Mava, trabeculae were subjected to constant velocity stretches from 0.95 Lo to 1.175 Lo with stretch durations of 100s, 10s, 1s, and 0.1s. After each stretch, muscle length was held constant for 60s and then briefly reduced to 0.8 Lo to slacken the muscle and establish the zero force level. Between stretches, muscles were equilibrated for 60s at a length of 0.95 Lo. Such large magnitude stretches have been commonly used to probe the diastolic properties of cardiac muscle samples (15-17).

The relationship between muscle viscous force (Fη) and stretch velocity (V) was fit to the dose-response relation: Fη = Fη_min +_ V x (Fη_max -_ Fη_min)_ / (V_50 +_ V), where Fη_min is_ minimum value of Fη (in relaxing solution), and V_50 is_ the velocity resulting in a half maximal increase of Fη.

The relationship between Fη versus [Ca^2+^] was fit to the Hill equation: Fη = Fη_max x_ [Ca^2+^]*^n^*^H^ / ([Ca^2+^]*^n^*^H^ + EC_50*n*H_), where Fη_max is_ the maximum Ca^2+^-activated viscous force, EC_50 is_ the [Ca^2+^] at which Fη is 50% of Fη_max,_ and *n*H is the Hill coefficient reflecting the slope of the relationship at EC_50._

Finally, in a subset of 2 experiments, the relationship between developed force and [Ca^2+^] was compared to that of Fη versus [Ca^2+^]. Developed force in the absence of inhibitors was measured at the same muscle length (1.175 Lo) used to measure Fη.

### Sex differences

We compared stiffness properties of RV trabeculae from 10-week old adult male vs. female mice. For males, body weight was greater (23.9 ± 0.6 g, n=3), than for females (weight 20.6 ± 0.6 g, n=3, *P* = 0.017). Trabecula length (783 ± 72 μm, n=6), and width (154 ± 24 μm, n=6), was similar in males and females. There was a trend for muscle thickness to be greater in males (108 ± 7 μm, n=3 μm) than females (87 ± 4 μm, n=3, *P* = 0.056).

### Statistical analysis

Data are presented as mean ± SE. Statistical tests were performed using Prism 10 software (GraphPad Software, Inc., La Jolla, CA) with a significance level set at *P*<0.05.

## RESULTS

### Viscous force response to stretch

Figure 1 shows typical records of muscle length, sarcomere length and muscle force from a relaxed trabecula subjected to a large ramp stretch of 10 s duration. Muscle length was linearly increased from 0.95 Lo to 1.175 Lo, associated with an increase of sarcomere length from ≈ 1.9 μm to 2.3 μm. In line with previous reports, stretch caused force to increase to a peak and then relax to a quasi-steady-state level after the stretch (6-9). This stress-relaxation did not involve an appreciable change in sarcomere length.

**Figure 1.**
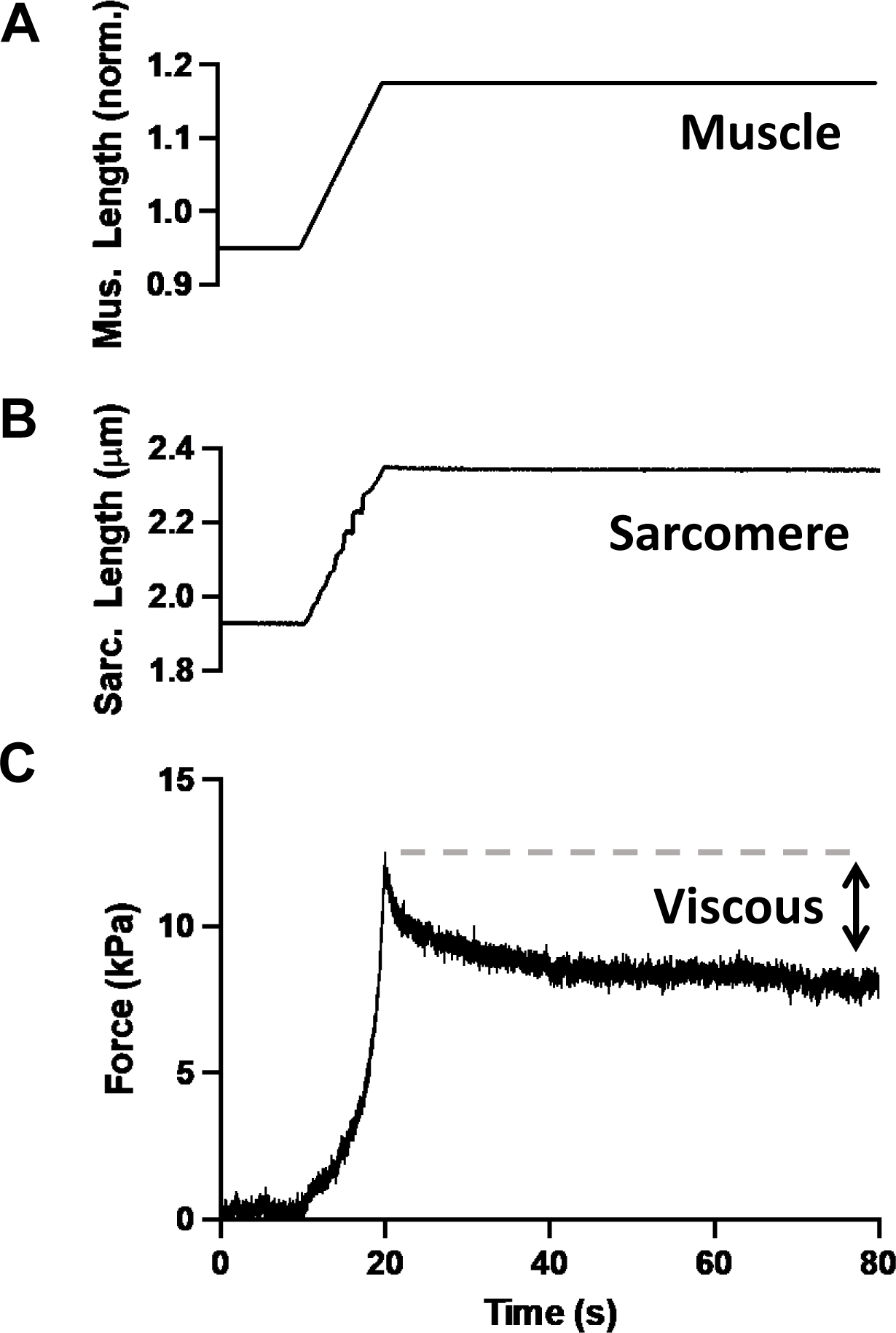
Viscous property of cardiac muscle. Example records of changes in muscle length, sarcomere length and muscle force in response to a linear muscle stretch under relaxing conditions. Viscous force of relaxed muscle was quantitated from the difference in the peak force at the end of the stretch versus the quasi-steady state level of force reached 60s after the stretch. Muscle length is normalized to a length at which the sarcomere length is 2.0 μm.

Peak force and the quasi-steady-state force level 60 s later were measured after muscle stretches with velocities varying over three orders of magnitude (Fig. 2A). Peak force and steady-state force were similar at the slowest stretch velocity (0.0025 muscle lengths per second (ML/s), stretch duration 100 s). However, as previously reported (6), the peak force after stretch progressively increased with increasing stretch velocity (Fig. 2A). In contrast, the steady-state force level (assessed 60s after stretch) did not appreciably change as a function of stretch velocity. Accordingly, the viscous force in response to stretch (defined as peak force minus steady-state force) was observed to increase with increasing stretch velocity in all experiments (Fig. 2B, P = 0.002 repeated measures one-way ANOVA). The dependence of the viscous force response on stretch speed was not linear.

**Figure 2.**
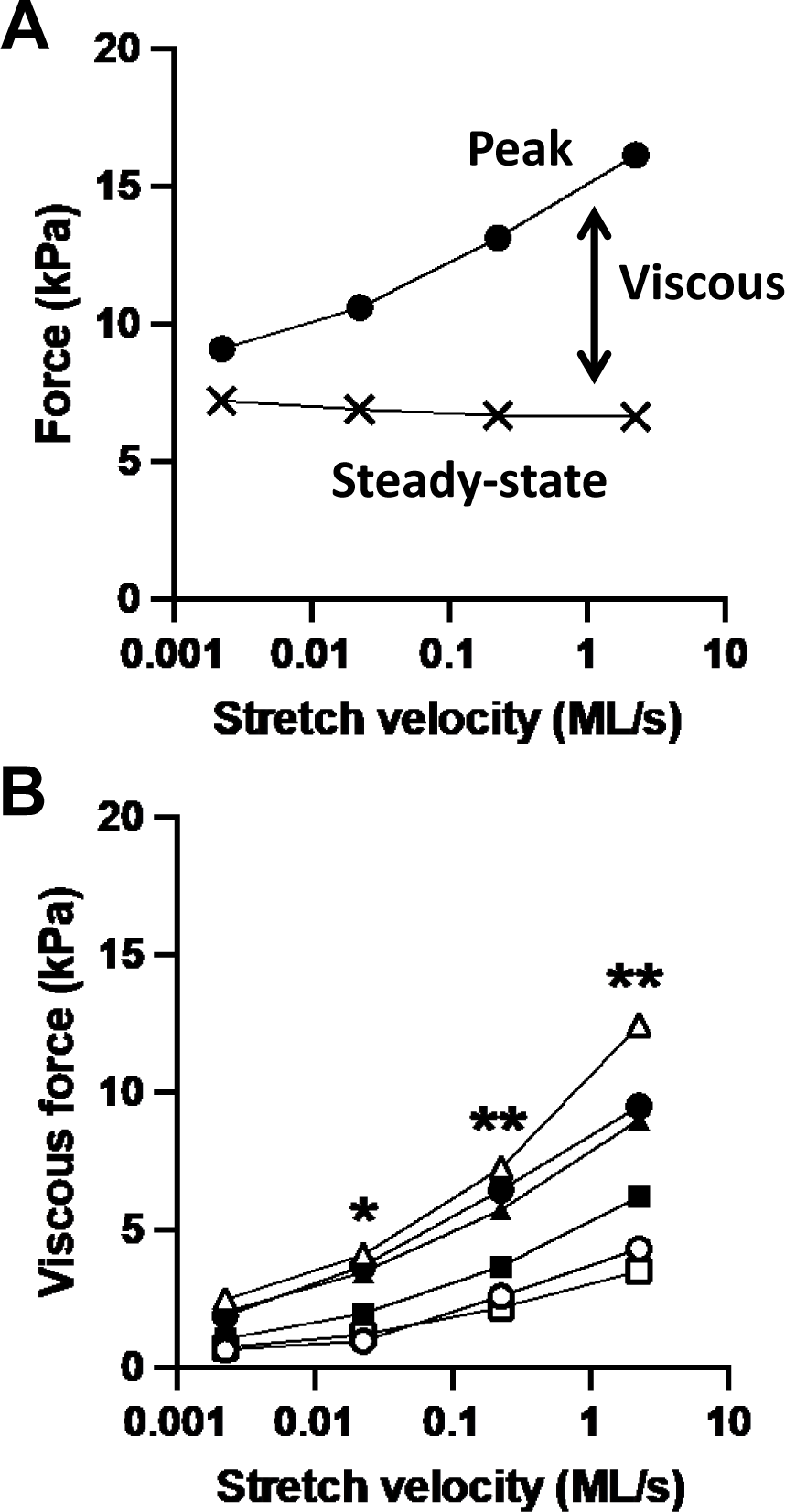
Velocity-sensitive viscous property. (A) Example data of peak and steady state force levels after stretches of relaxed muscle with velocities ranging over 3 orders of magnitude; note logarithmic abscissa scale for stretch velocity in muscle lengths per second (ML/s). (B) For all muscles in relaxing conditions, the viscous force (peak force minus steady-state force) increased with increasing stretch velocity (*P* = 0.002 repeated measures one-way ANOVA). Statistical comparisons are shown relative to the slowest stretch (* *P* <0.05, ** *P* < 0.01 with Bonferroni correction for multiple comparisons). For relaxed muscle, the effect of stretch on viscous force was not different for males (closed symbols) versus females (open symbols) (*P* = 0.57, repeated measures two-way ANOVA).

There was not a significant male vs. female difference observed in the effect of stretch velocity on viscous force (*P* = 0.57, repeated measures two-way ANOVA).

### Inhibition of force development by PNB plus Mava

Ca^2+^-activated force development was prevented after incubation of trabeculae with the cross-bridge inhibitors PNB (50 μM) plus Mava (50 μM) (Fig. 3A). In the presence of the inhibitors, muscle force measured in activating solution (0.9 ± 0.1 kPa, n = 6) was not increased relative to the passive force level measured in relaxing solution (0.95 ± 0.06 kPa, n = 6) (Fig. 3B).

**Figure 3.**
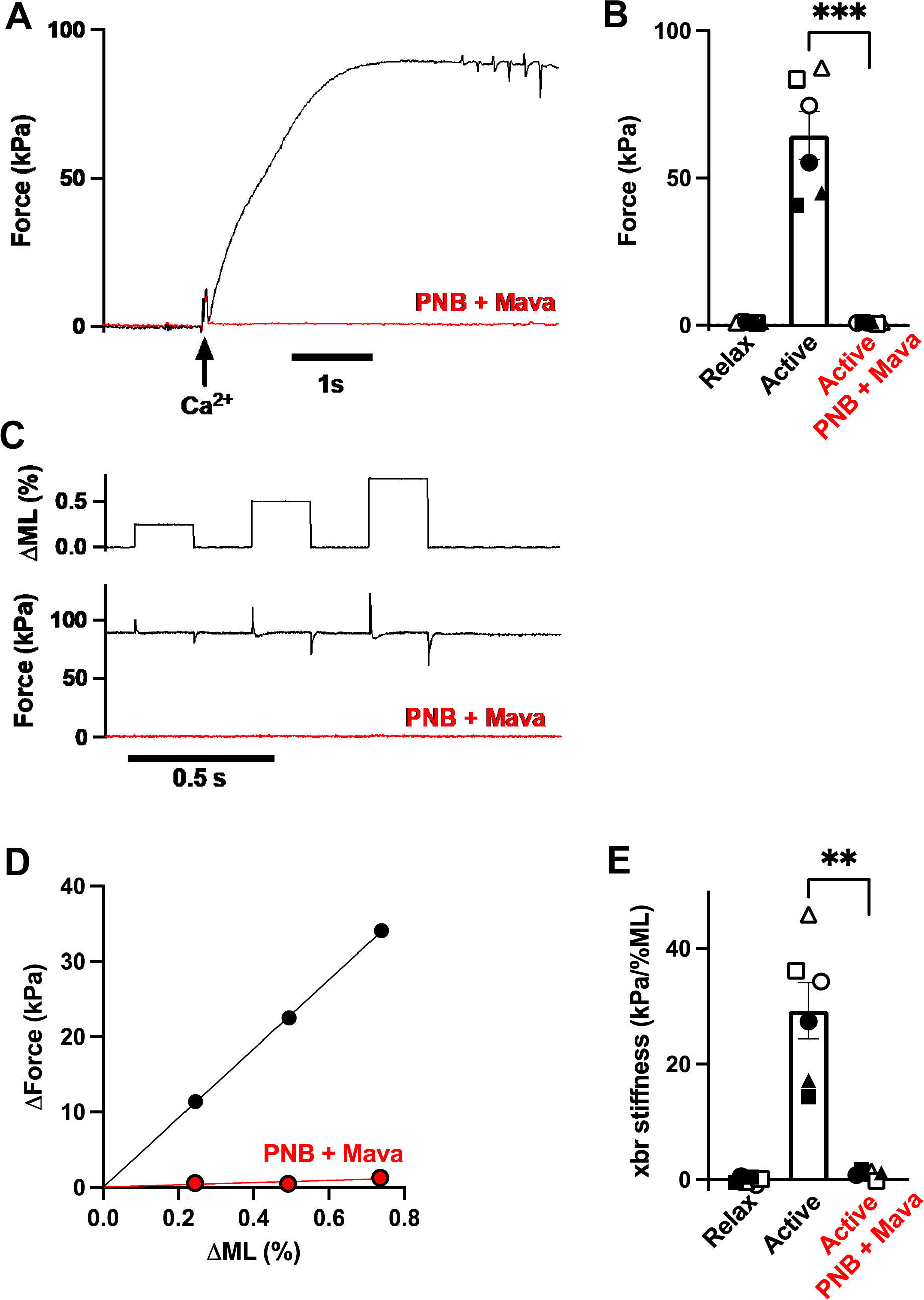
Inhibitors PNB + Mava eliminated force development. (A) Example records of muscle force response to Ca^2+^-activation (at arrow). Development of force (black trace) was completely inhibited in the presence of the inhibitors PNB plus Mava (red trace). (B) Summary of muscle force in the presence of low Ca^2+^ level (Relax), high Ca^2+^ level (Active), and high Ca^2+^ level with PNB + Mava completely inhibiting force development (*** *P* <0.001, paired t-test). Solid symbols show males, and open symbols show females. (C) Example records of muscle force measured at high Ca^2+^ level showing the effects of low amplitude (≤0.75%), rapid (1 ms), changes of muscle length (ΔML) both before and after addition of PNB + Mava. The presence of attached cross-bridges were evidenced from the force spikes induced by stretches. (D) Example data showing the relationship between the amplitude of the force spike versus the amplitude of the muscle stretch. The slope of this relationship was used as an index of muscle stiffness arising from attached cross-bridges. (E) Summary data of muscle stiffness due to attached cross-bridges in low Ca^2+^ (Relax), high Ca^2+^ (Active) and after addition of inhibitors eliminated attached cross-bridges (** *P* <0.01, paired t-test).

Prior to adding the inhibitors, maximum Ca^2+^-activated force was high, (64 ± 8 kPa, n=6) and interestingly, was higher in trabeculae from females (82 ± 4 kPa, n = 3) versus trabeculae from males (47 ± 4 kPa, n = 3, *P* = 0.004, unpaired t test) (Fig 3B).

To further test if the combination of PNB plus Mava inhibited muscle cross-bridges, we assessed the presence of attached cross-bridges using rapid (1 ms) small amplitude stretches (≤ 0.75% muscle length) to stress attached cross-bridges (Fig. 3C). In the absence of inhibitors, for activated muscle, attached cross-bridges were evidenced both from the high developed force and from transient increases of force in response to transient stretches (Fig. 3C). In contrast, with the inhibitors in the activating solution, elimination of attached cross-bridges was evidenced both from the absence of developed force, and from the lack of transient increases of force in response to transient stretches (Fig. 3C). Attached cross-bridges were quantitated from the slope of the relation between the amplitude of transient increases in force vs. the size of the transient stretch (Fig. 3D). In the presence of the inhibitors, the slope of this relation was close to zero with both high Ca^2+^ and low Ca^2+^, suggesting that the inhibitors fully prevented Ca^2+^-activation of cross-bridge attachment. However, in the absence of inhibitors, the slope of this relation was high, and was higher for trabeculae from females (39 ± 4 kPa/%ML, n = 3) versus males (20 ± 4 kPa/%ML, n = 3, *P* = 0.023, unpaired t test) (Fig. 3E), consistent with the higher contraction force noted for trabeculae from females (Fig. 3B).

In conclusion, under conditions of high Ca^2+^, the combination of inhibitors PNB plus Mava prevented the formation of load-bearing cross-bridges.

### Ca^2+^ increased viscous force response to stretch

Figure 4A shows typical records of changes in force in response to a 20% muscle stretch at the fastest stretch velocity (2.25 ML/s, duration 0.1 s) followed by a 60 s length hold. In the presence of the cross-bridge inhibitors, records were obtained, both in low Ca^2+^ (black trace) and then in high Ca^2+^ (red trace).

**Figure 4.**
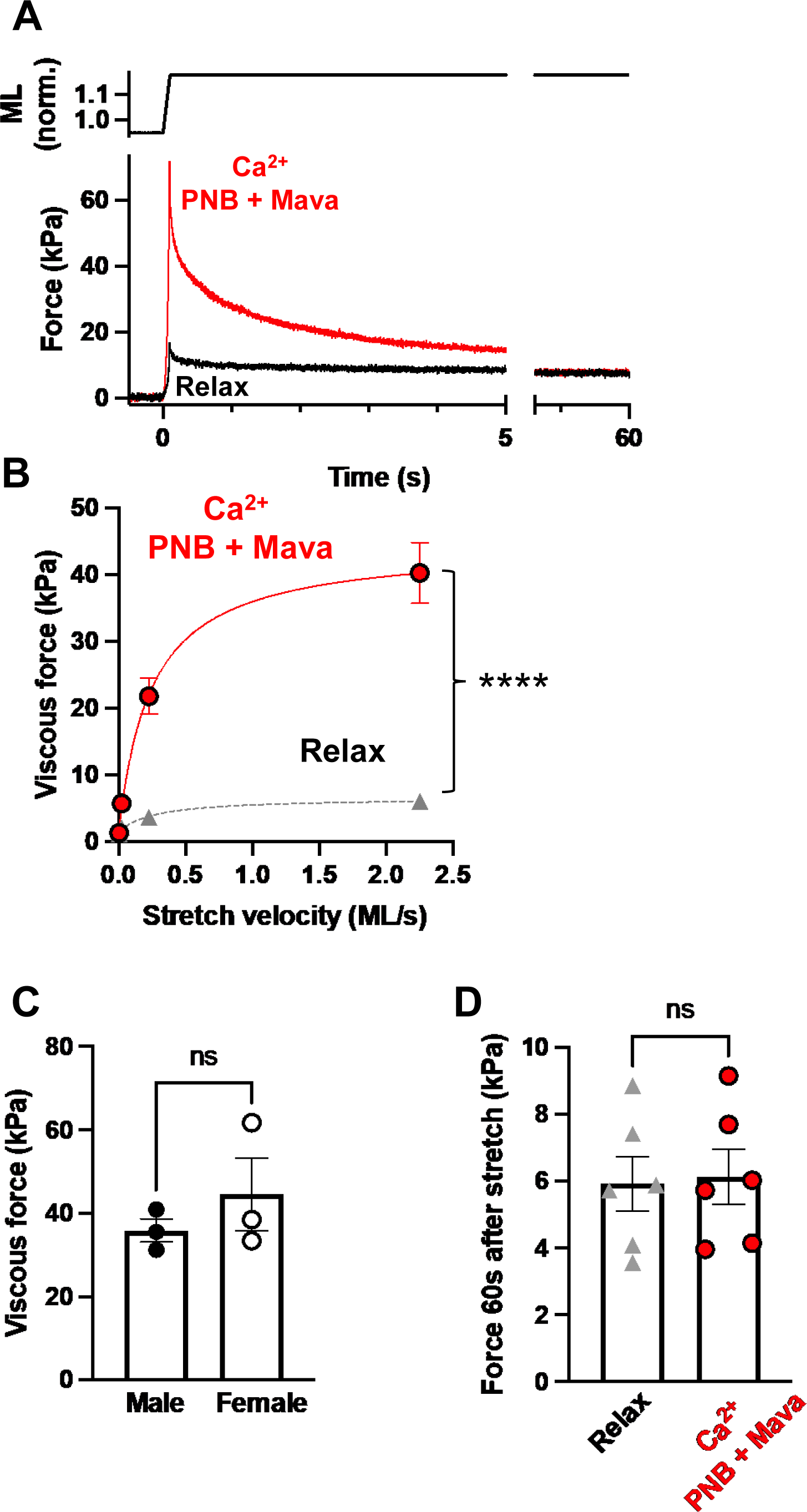
Viscous property is sensitive to Ca^2+^. (A) In the presence of cross-bridge inhibitors, example records showing that the peak force after a large amplitude muscle stretch was markedly increased by high Ca^2+^ level (red trace) compared to low Ca^2+^ level (Relax). (B) Relation of viscous force (peak minus steady state level) following stretch at various velocities and in the presence of cross-bridge inhibitors at both low Ca^2+^ (Relax), and high Ca^2+^ (**** *P* <0.0001 2-way ANOVA). (C) Viscous force response to the fastest stretch (velocity 2.25 ML/s) of trabeculae from males vs. females in the presence of cross-bridge inhibitors and high Ca^2+^. (D) Force level 60 s after 0.1 s stretch/hold protocol in the presence of cross-bridge inhibitors at both low Ca^2+^ (Relax), and high Ca^2+^.

In high Ca^2+^, there was no force developed prior to the stretch, indicating that the inhibitors prevented Ca^2+^-activated force development. However, with high Ca^2+^, the peak force after stretch was markedly increased compared to with low Ca^2+^. For all experiments, the viscous force response to stretch (peak force minus steady-state force 60s after stretch) was increased with stretch velocity (Fig. 4B) and was increased much more with high Ca^2+^ than with low Ca^2+^ (*P* < 0.0001, repeated measures two-way ANOVA). The extrapolated maximum with high Ca^2+^ (44.4 kPa) was > 6-fold higher than with low Ca^2+^ (6.6 kPa) (Fig. 4B). However, the estimated stretch velocity associated with a half-maximal increase in viscous force was similar in high Ca^2+^ (0.25 ML/s) versus low Ca^2+^ (0.26 ML/s) suggesting that the viscosity mechanism was similar at both high and low Ca^2+^. The observed dependence of force development on stretch velocity is explored in a theoretical model developed by Jezek et al. (18).

Despite the higher developed force noted for trabeculae from females versus males, there was no sex difference in the viscous force response to stretch (Fig. 4C).

The force measured 60s after stretch was not increased by Ca^2+^ (Figs. 4A & 4D). This finding is consistent with the observed absence of force development in the presence of the inhibitors, and also suggests that there is no effect of Ca^2+^ on the muscle elastic property.

### Relationship of muscle viscous property to Ca^2+^ level

The dependence of the viscous force component on Ca^2+^ level was assessed using the response to the fastest stretch velocity (2.25 ML/s) over a range of Ca^2+^ levels. The viscous force vs. pCa relation (Fig. 5) had a sigmoidal form that was well described by the Hill equation (EC_50_ = 1.3 μM, *n*H = 2.5, n = 6). The value for relaxed muscle was 16% of the value for maximal Ca^2+^-activation. This relationship did not differ between males and females (*P* = 0.45, 2-way repeated measures ANOVA).

**Figure 5.**
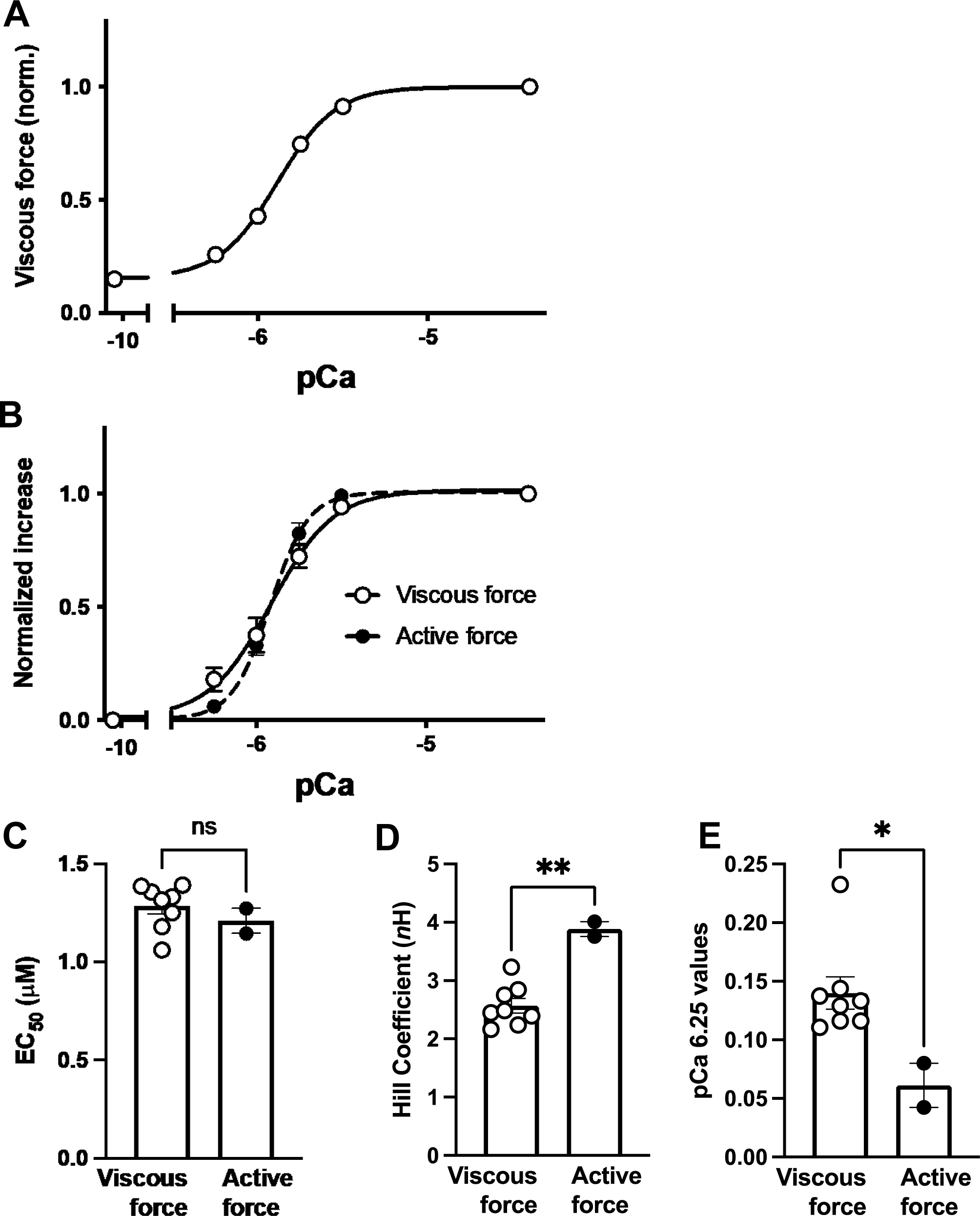
Ca^2+^ regulation of viscous property. (A) Effect of Ca^2+^ level (in the presence of cross-bridge inhibitors) on the viscous force response to the fastest stretch (velocity 2.25 ML/s) (mean values, n=6, error bars smaller than symbols). Data were fit to the Hill equation. (B) In 2 additional experiments, the viscous force-pCa relation in the presence of inhibitors is compared to the developed force-pCa relation in the absence of inhibitors (means ± se). (C&D) Values from fit to the Hill equation for EC_50_ and slope. (E) Values measured at pCa 6.25. *P < 0.05, **P<0.01 paired t-test.

In a subset of experiments (n=2), the effect of Ca^2+^ on active force development (in the absence of inhibitors) was directly compared in the same muscles to the effect of Ca^2+^ on the viscous force response to stretch (in the presence of the inhibitors) (Fig 5 B). To minimize length-dependent effects on Ca^2+^-activation, both force development and the viscous force response to stretch were measured at the same muscle length (1.175 Lo). The viscous force-pCa relation was similar to the developed force-pCa relation (Fig. 5B) and the corresponding EC50 values were not different (Fig. 5C). The viscous force-pCa relation had a shallower slope, as reflected in a lower Hill Coefficient (Fig. 5D). Accordingly, at pCa 6.25, the relative increase of viscous force was greater than the increase of developed force (Fig. 5E).

### Temperature-independent muscle viscous property

As a further check for a contribution of cross-bridge activity to the increased viscous force response to stretch at high Ca^2+^ level, we assessed the effect of temperature. Reducing the temperature from 21°C to 10°C did not reduce the viscous force response to stretch assessed at the fastest stretch velocity (Fig. 6A).

**Figure 6.**
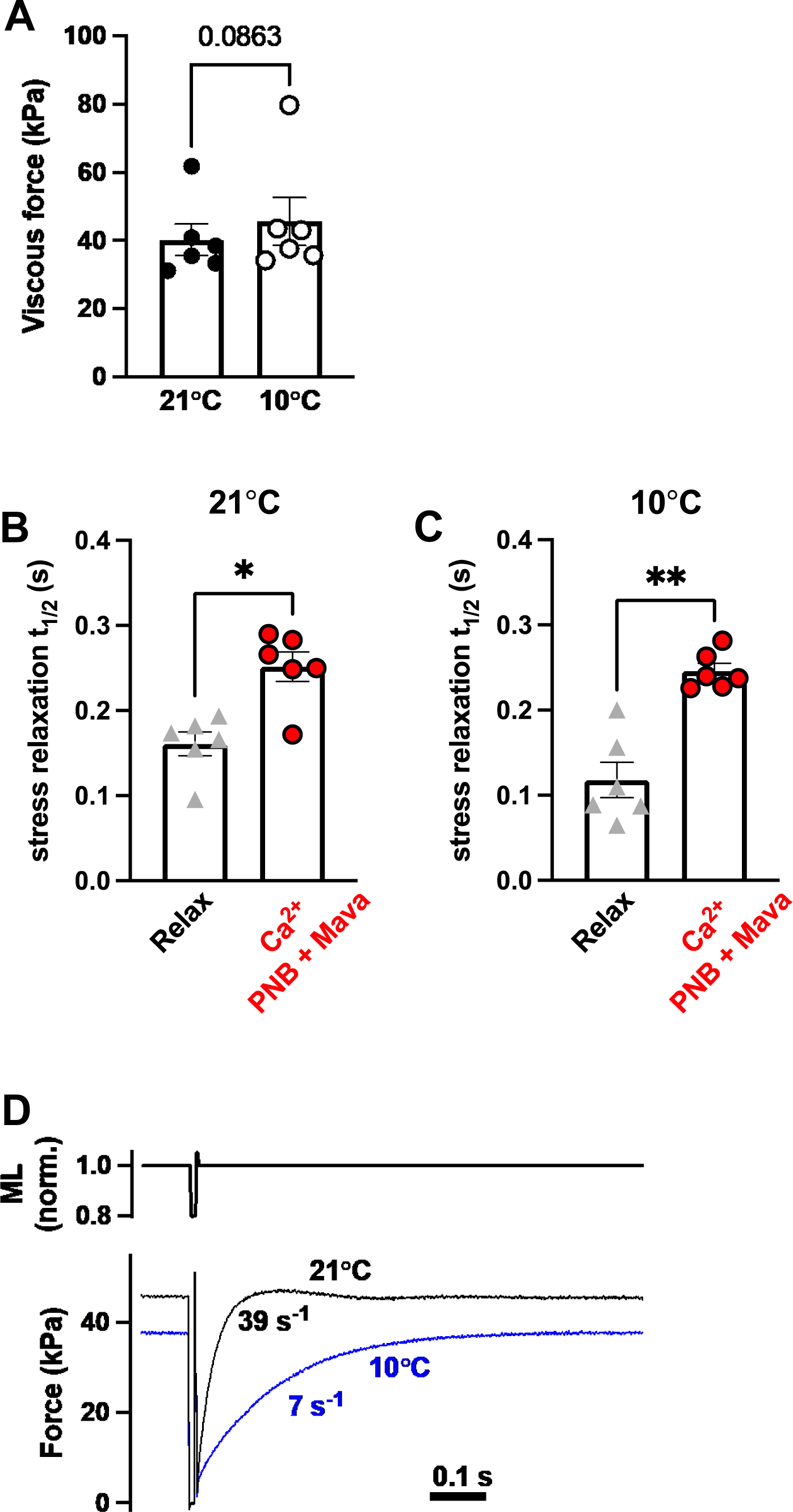
No effect of temperature on Ca^2+^-activated viscous property. (A) Ca^2+^-activated viscous force is not decreased by lowering temperature from 21°C to 10°C. (B&C) Stress relaxation half time (t_½)_ measured directly from records for trabeculae in the presence of inhibitors and both low Ca^2+^ (Relax) and high Ca^2+^, and with temperature set first to 10°C and then 21°C. (D) Low temperature reduced developed force and slowed the rate constant of force redevelopment after cross-bridges were mechanically disrupted.

We also assessed the effect of temperature on the time course of stress relaxation after a stretch. The half time of stress relaxation (t_½)_ with high Ca^2+^ level was significantly higher compared to with low Ca^2+^ level (Fig. 6B). However, t_½_ was not affected by reducing the temperature to 10°C. In contrast, cross-bridge activity is known to be impaired with low temperature (19). Therefore, a lack of sensitivity to temperature is consistent with the stress relaxation process not involving cross-bridge activity. To evaluate the effect of temperature on cross-bridge activity in our preparation, for one experiment where force was measured in the absence of inhibitor, as expected the maximum force level was lower at 10°C than 21°C (37 vs 46 kPa) (Fig. 6D). Moreover, the rate constant for force redevelopment after mechanically disrupting cross-bridges was appreciably slower at 10°C than 21°C (Fig. 6D). To minimize a potential impact from deterioration of the preparation with multiple activations, we first studied contraction at 10°C and then at 21°C.

In summary, viscous force is not sensitive to temperature, but actively developed force is sensitive to temperature. This difference suggests that different mechanisms are involved in force development versus viscous force and further suggests that cross-bridge cycling does not contribute to the Ca^2+^-sensitive viscous property.

Finally, in the absence of inhibitors, the rate constant for force redevelopment after mechanically disrupting cross-bridges was 35 ± 5 s^-1^, n = 6, corresponding to a t_½_ of 0.02 ± 0.005 s. Thus, the time course of force redevelopment is approximately an order of magnitude faster than the time course of stress relaxation (Fig. 6B), reinforcing the idea that force development and muscle viscosity involve different underlying phenomena.

### Ca^2+^ effect on muscle viscous property requires intact myofilaments

For trabeculae subjected to a high-salt extraction protocol designed to remove the myofilaments, the viscous force measured after the most rapid stretch was reduced (Fig. 7) from 6.1 ± 0.9 kPa to 3.9 ± 0.7 kPa, n= 6. This suggests that at the extended muscle length attained after a large 20% stretch, the viscous force measured after the stretch is partly attributed to myofilament-based structures and party to non-myofilament structures, such as the extracellular matrix. The relative contributions of forces associated with myofilament versus non-myofilament structures is addressed in the theoretical study by Jezek et al. (18).

**Figure 7.**
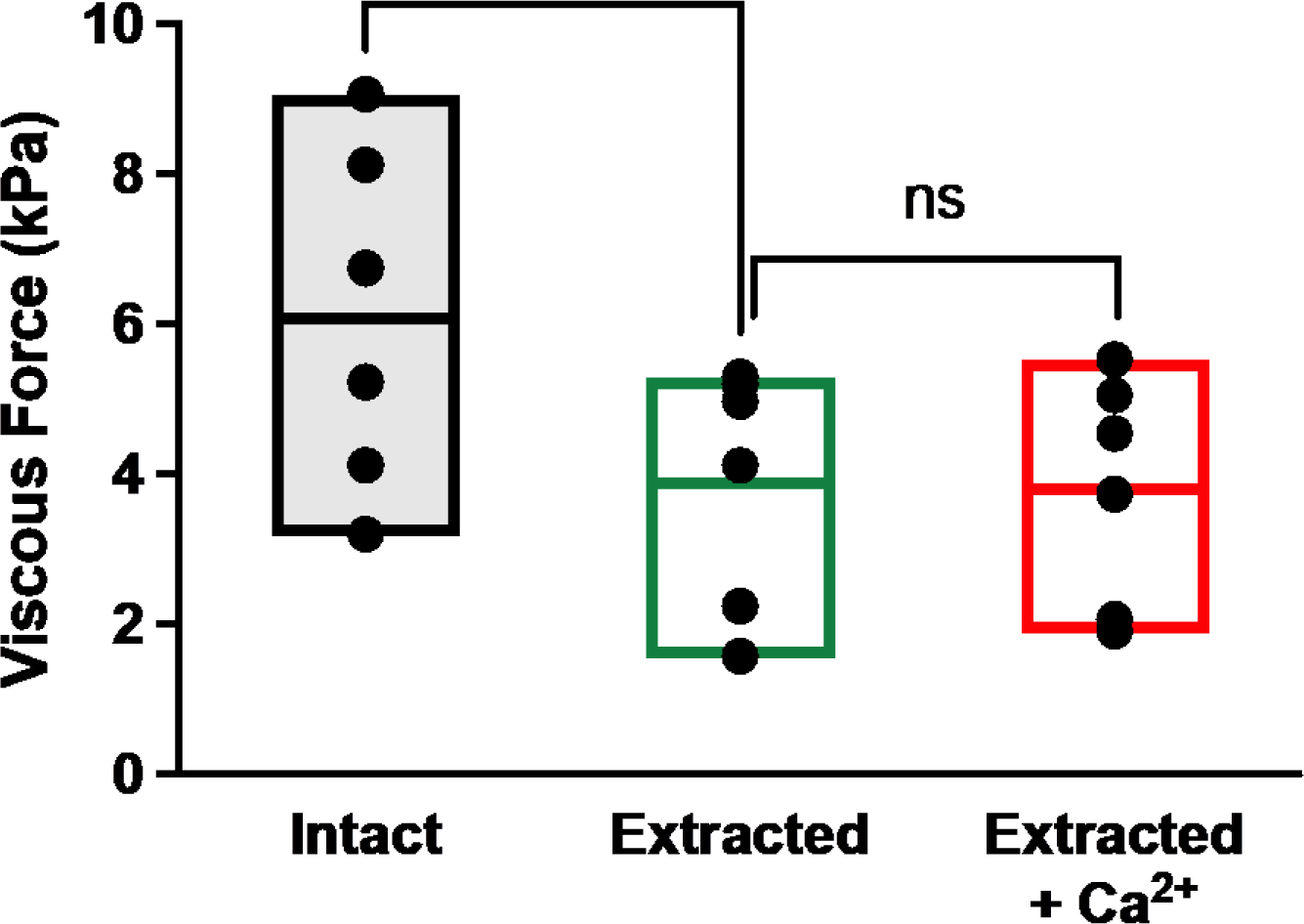
Effect of Ca^2+^ to increase viscous force was lost after extraction of myofilaments. Viscous force measured in the presence of inhibitors at low- and high Ca^2+^ level for trabeculae before (intact) and after high-salt extraction of the myofilaments (mean and high and low values, **P<0.01 paired t-test).

Importantly, after myofilament extraction, the effect of Ca^2+^ to increase the viscous response to muscle stretch was eliminated (Fig. 7) indicating that the large Ca^2+^-dependent component of the apparent muscle viscosity is entirely a myofilament property.

## DISCUSSION

The major finding of this study is that Ca^2+^ regulates a viscous property of cardiac muscle independent of an effect of Ca^2+^ on cross-bridge cycling. This phenomenon considerably expands the role of Ca^2+^ for impacting cardiac muscle mechanical properties where Ca^2+^ mediates both activation of contraction and an increase in muscle stiffness. This effect of Ca^2+^ on a muscle viscous property might influence the mechanical response of myocardium to fluctuating Ca^2+^ levels during activation and relaxation. Building on our observations, Jezek et al. (18) have developed a theoretical model to explore the potential effects of Ca^2+^ on the passive mechanical properties of the myocardium.

### Viscous property of relaxed muscle

We observed the familiar biphasic stress response to a ramp increase in muscle length. Force increased during the stretch and reached a peak at the completion of the stretch. Following the ramp stretch, when the muscle length was held at a constant final length, the stress decayed to a steady level. Stress-relaxation of cardiac muscle (6-9), is consistent with a viscoelastic material response.

The observed peak stress increased with increasing stretch velocity, even as the final length was the same for all ramp speeds. This dependence of peak stress on the stretch velocity is consistent with a nonlinear viscous component of the passive mechanical response of the muscle to stretch (6). Importantly, the magnitude of the viscous response was sensitive to stretch velocity over a physiologically relevant range of stretch velocities, suggesting relevance to the systolic and diastolic function of the beating heart.

Multiple structural elements contribute to cardiac muscle viscous properties; with approximately equal contributions reported to arise from titin, actin, microtubules and intermediate filaments (2). For muscles subjected to high stains, such as the stretch protocol used in this study, then a significant contribution also arises from the extracellular matrix (2).

In an accompanying paper we present a theoretical analysis of the observed stress response to stretch that suggests that the passive stress relaxation proceeds over a power-law time course that is governed in part by the molecular mechanics of the elastic domain of titin in the I band of the sarcomere (18).

### Inhibition of cross-bridge cycling

Characterizing an effect of Ca^2+^ on the muscle viscous property relies crucially on effectively inhibiting Ca^2+^-activation of cross-bridge cycling. Preliminary studies revealed that 50 μM PNB alone did not completely inhibit Ca^2+^-activated force development (residual force was ∼ 10%). Therefore, to completely inhibit force development, the current study combined two inhibitors: 50 μM PNB plus 50 μM Mava.

Several lines of evidence indicate that effective inhibition of contraction was achieved by the combination of PNB plus Mava. Specifically, in the presence of the inhibitors and a high Ca^2+^ level: a) Ca^2+^-mediated force was abolished; b) attached cross-bridges were eliminated (assessed by rapid stretches of small amplitude); and c) the force level 60s after stretch was not increased relative to relaxed muscle, indicating no active force generation. Moreover, additional evidence excludes effects due to cross-bridge cycling in the effect of Ca^2+^ on the muscle viscous property. In the presence of the inhibitors and high Ca^2+^ level: a) although low temperature impaired cross-bridge cycling, low temperature did not reduce the viscous force or slow the time course of viscous force relaxation; and b) the time course of viscous force relaxation was an order of magnitude slower than expected for a cross-bridge-mediated effect.

### Ca^2+^ activation of cardiac muscle viscous stiffness

Recent studies reported that electrical activation of frog skeletal muscle increased the passive component of stiffness after active force development was prevented by the cross-bridge inhibitor PNB (1). An increase in intracellular Ca^2+^ level following electrical stimulation was suggested to increase muscle stiffness mediated by the structural protein titin. Indeed, previous studies have suggested Ca^2+^-mediated stiffening of titin involves Ca^2+^ binding to the PEVK region (20) or to the immunoglobulin domain (21).

The current study directly tested the effect of activator Ca^2+^ level on cardiac muscle stiffness. In the presence of the cross-bridge inhibitors PNB plus Mava, a high Ca^2+^ level increased the magnitude of the viscous force response to muscle stretch by ≈ 6-fold relative to the relaxed state. The dependence of viscous stiffness on the level of activator Ca^2+^ followed a sigmoidal relationship with the same sensitivity to Ca^2+^ as observed for Ca^2+^ activation of myofilament force development. This suggests a common regulatory mechanism resulting in a coordinated effect of Ca^2+^ to both increase force development and increase viscous stiffness. Nevertheless, at pCa 6.25 the effect of Ca^2+^ on the viscous property was greater compared to the effect of Ca^2+^ on force development. Possibly, for low levels of Ca^2+^ activation in-vivo, Ca^2+^ might increase viscosity more than contraction.

The effect of Ca^2+^ to increase the muscle viscous property was abolished after a high salt protocol to extract the myofilaments, indicating that the Ca^2+^ effect involves a myofilament-based mechanism. In contrast, as noted above, for low Ca^2+^ level, the viscous property depends on both myofilament and non-myofilament structures. Interestingly, despite the large increase in viscous force with high-versus low Ca^2+^ level, the dependence of viscous force on stretch velocity was similar for both high- and low Ca^2+^ levels. This suggests that there is matching of the myofilament-based and non-myofilament-based components of the muscle of viscous property.

The mechanism for the effect of Ca^2+^ to cause a large increase in the muscle viscous property was not determined in this study. Previously, interaction between actin and titin was proposed to increase muscle stiffness (22-24). Moreover, recent studies suggested that with electrical stimulation of frog skeletal muscle increased stiffness resulted from switching of titin from an extensible spring (OFF-state) to a mechanical rectifier (ON-state) that allows shortening but has an elevated resistance to stretch (1). The titin ON state and increased muscle stiffness was hypothesized to involve cross-linking of flexible titin to relatively stiffer actin filaments (1). For this study, we used a computational model to quantitatively investigate Ca^2+^-dependent binding of titin to actin and show that this mechanism can account for both effects of Ca^2+^ on stiffness and effects on the time-course of stress relaxation (18).

### Sex difference

Trabeculae from both males and females were tested. Muscle viscous and elastic properties were not different in males vs. females. Interestingly, trabeculae from females developed significantly higher forces than males. Consistent with higher developed force, estimates of attached cross-bridges using small rapid stretches were higher in females than males. The reason for this difference is not clear. Trabeculae from females tended to be thinner, which might result in some overestimation of force development. Further studies of this sex difference in force development are warranted.

## Conclusion

In addition to Ca^2+^-activation of contraction, Ca^2+^ also markedly increased the apparent viscous force response to rapid stretch of cardiac muscle. This behavior opens a new window on Ca^2+^ modulational of cardiac muscle mechanical properties that may have implications for understanding both systolic and diastolic properties.

## ACKNOWLEDGEMENTS

This work was supported by Department of Veterans Affairs Merit Review Award I01BX000740 (AJB) and National Heart, Lung and Blood Institute Grant R01 HL154624 (A.J.B., D.A.B., N.C.C).

## Disclosures

None

## Notes

### Competing Interest Statement

The authors have declared no competing interest.

